# Clinical Relevant Immunosuppressive Drugs Differentially Modulate Axonal Outgrowth from Human Stem Cell–Derived Neurons

**DOI:** 10.64898/2026.06.29.735084

**Authors:** Gunnar H.D. Poplawski, C Weinholtz, G Woodruff, R Ahmad, W Bunner, Rodrigo Gonzales, MH Tuszynski

## Abstract

Neural stem cell (NSC) transplantation is a promising strategy for repairing the injured spinal cord, but transplanted cells typically require immunosuppressive therapy to prevent rejection—even for induced pluripotent stem cell (iPSC)-derived autologous grafts. However, the effects of immunosuppressive drugs on neurite outgrowth and axonal regeneration— processes critical for neural circuit reconstruction—have not been fully characterized. In this study, we tested nine clinically relevant immunosuppressants on human iPSC-derived neurons and primary human spinal cord NSCs in vitro at concentrations approximating clinical exposure levels. The drug panel included FK-506 (tacrolimus), cyclosporine A (CsA), rapamycin, belatacept (Nulojix), etanercept (Enbrel), mycophenolate mofetil (CellCept), cyclophosphamide (Cytoxan), prednisone, and azathioprine (Imuran). Neurite outgrowth was quantified via automated high-content imaging. Multiple agents, including CsA, Imuran, Nulojix, and CellCept, induced significant reductions in neurite outgrowth in a cell type- and dose-dependent manner, with **CsA producing the most robust and consistent inhibition** across both cell lines. In contrast, FK-506 showed no significant effect on neurite extension at clinically relevant concentrations. Consistent with the in vitro results, human neural progenitor cell grafts in a rodent spinal cord injury model exhibited significantly reduced graft-derived axon extension in the host spinal cord when hosts were treated with CsA rather than FK-506. These findings demonstrate that immunosuppressant choice can profoundly influence neural graft integration and axonal regeneration. Our study underscores the importance of preclinical evaluation of immunosuppressive regimens and suggests that selecting agents such as FK-506 over CsA may improve outcomes in future stem cell–based therapeutic trials for spinal cord injury and related disorders of the central nervous system.

**Highlights:**

- Cyclosporine A inhibits axon outgrowth in human neurons in vitro
- FK-506 preserves neurite extension across stem cell–derived neuron types
- Axon outgrowth is reduced in vivo with CsA but not FK-506
- Immunosuppressant selection critically affects neural graft integration
- FK-506 may be preferable to CsA for SCI cell transplantation protocols

## INTRODUCTION

Spinal cord injury (SCI) is a devastating condition that often results in permanent neurological deficits due to extensive cell loss and the failure of damaged axons to regenerate. After the primary trauma, a cascade of secondary injury mechanisms — including inflammation, excitotoxicity, and glial-scar formation—further impedes recovery^1^. Spontaneous repair in the adult spinal cord is extremely limited, so intensive research has turned toward regenerative therapies to restore lost function. In particular, cell-based strategies—such as grafts of neural stem or progenitor cells—have shown promise in replacing lost neurons and glia, rebuilding circuitry, and promoting recovery in animal models^2,3^. Transplantation of exogenous neural progenitor cells (NPCs) can provide a cellular substrate for regeneration, secrete neurotrophic factors, and remyelinate axons, thereby helping to reconstruct injured neural networks^2,4-6^.

These encouraging pre-clinical findings have laid the groundwork for translating stem-cell therapies to the clinical setting of SCI. Over the past decade, multiple early-phase clinical trials have explored the safety and potential efficacy of stem-cell grafts in SCI patients. For example, human embryonic stem-cell (hESC)-derived oligodendrocyte progenitor cells (OPCs) were transplanted into individuals with subacute SCI in the Geron/Asterias “SCiStar” trial. This approach proved feasible and safe, with reports that most participants experienced some neurological improvement (e.g., recovery of one or more levels of sensory or motor function) following the OPC injections^7^.

In another Phase I trial, researchers implanted allogeneic fetal spinal cord–derived neural stem cells (NSCs; NSI-566 cell line) into patients with chronic SCI^8^. After 18–27 months of follow-up, no serious adverse effects were noted, and two of the four treated patients showed detectable gains in neurological function (improvement of one to two levels on the International Standards for Neurological Classification of SCI)^8^. These results, while preliminary, demonstrate the potential of stem-cell transplantation to partially improve outcomes even in chronic, established SCI. Building on extensive pre-clinical work by several groups, the first-in-human trial of induced pluripotent stem-cell (iPSC)-derived neural progenitors for subacute SCI has recently been initiated in Japan^9^. Together, these efforts reflect a growing consensus that stem-cell-based interventions may complement traditional approaches and address the unmet need for restoring function after SCI.

A major challenge for any cell-transplantation strategy is overcoming immune rejection of the graft. Autologous iPSC-derived cells were originally proposed as an immune-evasive solution, since they genetically match the host. However, contrary to early assumptions, studies have shown that even autologous or syngeneic iPSC grafts can elicit immune responses. In a seminal study, Zhao et al. found that syngeneic mouse iPSC derivatives formed teratomas that were infiltrated by T cells and ultimately rejected, despite the cells being “self” to the host^10^. Although subsequent reports have offered mixed findings—for instance, other groups observed minimal immune reactions to autologous iPSC-derived cells in certain contexts^11^—the risk of immunogenicity remains a significant concern. In the clinical setting, nearly all stem-cell transplantation protocols for SCI (even those using cells presumed to be patient-compatible) have included some form of immunosuppressive therapy to promote graft survival^12^. For example, the ongoing iPSC-NPC trial in Japan is administering transient immunosuppression to patients receiving the grafts^9^. Likewise, in prior human trials of allogeneic NSC transplantation, patients were placed on immunosuppressant regimens and showed no signs of graft rejection or anti-donor immune responses during the treatment period^8^. These observations underscore that, despite the potential for immune matching with iPSC technology, effective immunosuppression is still usually required to protect neural grafts from host immune attack.

Current immunosuppressive regimens for cell transplantation in the central nervous system are largely adapted from those used in organ-transplant medicine. Calcineurin inhibitors such as cyclosporine A (CsA) and tacrolimus (FK-506) are among the most widely used agents; they prevent T-lymphocyte activation by blocking the calcineurin–NFAT signaling pathway required for interleukin-2 (IL-2) production. In the context of experimental SCI therapies, calcineurin inhibitors have been extensively employed to prevent rejection of engrafted neural stem cells. For instance, many rodent studies of human NSC grafts rely on daily CsA administration to ensure long-term cell survival^13^. Tacrolimus has also been used in human SCI trials: in the Asterias/Lineage OPC1 trial, patients received low-dose tacrolimus for about 60 days post-transplantation as a precaution against immune rejection^7^.

Another important class of immunosuppressants are the mTOR inhibitors (e.g., rapamycin/sirolimus), which act by inhibiting IL-2-driven T-cell proliferation. These are sometimes used as alternatives or adjuncts to calcineurin inhibitors in transplantation protocols, owing to their distinct mechanism of action and potential to promote tolerance. Other immunosuppressive approaches can include anti-proliferative drugs like mycophenolate mofetil or azathioprine (which impair lymphocyte DNA synthesis), corticosteroids (which broadly dampen immune and inflammatory responses), and monoclonal antibodies such as basiliximab (which blocks the IL-2 receptor on T cells).

Indeed, multi-modal immunosuppressive regimens have been tested in SCI cell-therapy trials—for example, one Phase I study combined an IL-2R antibody (basiliximab) with tacrolimus and mycophenolate for 12 weeks in order to prevent rejection of an NSC graft^14^. Each of these immunosuppressive agents carries a unique profile of action and side effects, and understanding their impact in the context of neural stem-cell engraftment is clinically relevant.

While immunosuppression is clearly effective at improving graft survival, much less is known about how these drugs influence the regenerative potential of neural stem-cell grafts— particularly the outgrowth of axons and the integration of graft-derived neurons into host circuits. Axonal extension from transplanted neurons (and promotion of host-axon sprouting) is thought to be a key contributor to functional recovery, as new relay circuits must form across sites of SCI damage for meaningful neurological improvement^14^. However, the potential interactions between immunosuppressant drugs and axon-growth mechanisms have not been thoroughly investigated. On one hand, immunosuppressants might aid regeneration indirectly by reducing graft inflammation and glial scarring. On the other hand, they could also have direct or indirect inhibitory effects on neurons: for example, systemic calcineurin inhibition could conceivably alter axon-guidance signaling, and mTOR inhibition is well known to reduce intrinsic axon-regenerative capacity in some contexts (since active mTOR pathways are associated with robust axon growth^15^). To date, only limited data address these possibilities. Notably, one prior study reported that common immunosuppressants can affect human NSC biology in vitro (e.g., influencing cell proliferation and differentiation), but found little apparent impact on graft outcomes in vivo in an SCI model^16^.

This paradox highlights our incomplete understanding: even as immunosuppressive therapy becomes routine in cell transplantation, its consequences for neural-repair processes like axonal regeneration remain uncertain. In short, beyond preventing immune rejection, do these drugs hinder, help, or have no effect on the ability of grafted neurons to extend axons and reconnect neural pathways? This critical question has yet to be fully answered.

Given this gap in knowledge, the present study was designed to systematically examine how clinically relevant immunosuppressants influence axon outgrowth from human neural stem-cell grafts, both in vitro and in vivo. We focused on several immunosuppressive agents that are commonly used or proposed in transplantation protocols—including a calcineurin inhibitor (FK-506/tacrolimus), an mTOR inhibitor (rapamycin), and others—and evaluated their effects on neurite extension from human stem-cell-derived neurons. In cultured human neural stem/progenitor cells (derived from hiPSCs and from fetal spinal cord tissue), we assessed axonal outgrowth in the presence of these drugs to simulate the in vitro environment under immunosuppressive conditions. Additionally, using a rodent model, we transplanted human fetal NSCs into the spinal cord and treated the hosts with FK-506/tacrolimus or rapamycin, then measured the extent of axonal elongation from the grafts within the injured spinal cord. Our goal was to determine whether these commonly used immunosuppressants modulate the growth and integration of neural stem-cell grafts. By clarifying the neurobiological impacts of immunosuppressive therapy, this work aims to inform future transplantation strategies for SCI—balancing the need for immune protection of grafts with the ultimate goal of maximizing neuronal regeneration and functional recovery.

## MATERIALS AND METHODS

### Drug Preparation and Dosing

Immunosuppressive agents included FK-506 (tacrolimus), cyclosporine A (CsA), rapamycin, belatacept (Nulojix), etanercept (Enbrel), mycophenolate mofetil (CellCept), cyclophosphamide (Cytoxan), prednisone, and azathioprine (Imuran). Stock solutions were prepared at 25 or 50 mg/ml in sterile DMSO or water and diluted into culture media to final concentrations ranging from 0.01 to 10 µg/ml. Drugs were sourced from Sigma-Aldrich, Selleck Chemicals, or commercial pharmaceutical suppliers. All compounds were added to neuronal cultures 2 hours after plating.

### Immunocytochemistry and Neurite Outgrowth Quantification

Cells were fixed in 4% paraformaldehyde (PFA) in PBS for 20 minutes at room temperature, permeabilized, and blocked with PBS containing 0.25% Triton X-100 and 5% donkey serum for 1 hour. Cells were incubated with mouse anti-βIII-tubulin primary antibody (TUJ1; 1:2000, Promega) for 2 hours at room temperature, followed by Alexa Fluor 568-conjugated anti-mouse secondary antibody (1:1000, Life Technologies) and DAPI (1 µg/ml) for 1 hour. After washing, images were acquired at 4× magnification using an ImageXpress Micro system (Molecular Devices).

Neurite outgrowth was automatically quantified using the MetaXpress neurite outgrowth module (Molecular Devices). Values were normalized to untreated PDL control wells to account for experiment-to-experiment variability and are presented as mean ± SEM.

### Human iPSC Generation and Differentiation

Human iPSCs were generated from patient-derived fibroblasts and differentiated into neurons as described^17^. Briefly, TRA1-81+ hiPSCs were co-cultured with PA6 stromal cells to induce neural differentiation. On day 11, cells were dissociated with Accutase, and NPCs were purified by FACS using CD184+CD15+CD44−CD271− gating. NPCs were expanded in the presence of bFGF.

Following 3 weeks of neuronal differentiation (bFGF withdrawal), CD24+CD184−CD44− neurons were isolated via FACS and plated in 384-well plates (Matrix, Thermo Fisher) at 2,500 cells/well in 50 µl media containing DMEM/F12 + GlutaMAX, 0.5× N2, 0.5× B27, 1× penicillin/streptomycin, 0.5 mM dbcAMP (Sigma), 20 ng/ml BDNF, and 20 ng/ml GDNF (Peprotech). Plates were pre-coated with poly-ornithine (40 µg/ml, Sigma-Aldrich) and laminin (5 µg/ml).

### Human Fetal Spinal Cord–Derived NPC Culture

The human spinal cord–derived NPC line UCSD 1113, isolated from a 9-week-old embryo^18^, was maintained on CELLstart-coated plates in a 1:1 mixture of DMEM/F12 and Neurobasal medium supplemented with 1× N2, 1× B27, 2 mM GlutaMAX, 10 ng/ml human LIF (Stemgent), and 20 ng/ml each of FGF2 and EGF (Peprotech). For terminal differentiation, media were switched to differentiation medium containing 300 ng/ml cAMP and 0.2 mM vitamin C for 7 days. Differentiated cells were dissociated with Accumax and replated on PDL-coated 48-well plates (20 µg/ml; Sigma-Aldrich).

### In Vivo Transplantation of Human NPCs

All animal procedures were conducted in accordance with protocols approved by the Institutional Animal Care and Use Committee (IACUC) at the University of California, San Diego. Female athymic nude mice (8–10 weeks old) were anesthetized with ketamine (100 mg/kg) and xylazine (10 mg/kg) and placed in a stereotaxic frame. A C6 laminectomy was performed, and 10 µl of GFP-labeled human fetal spinal cord–derived NPCs (2.5 million cells) were injected into the spinal cord parenchyma over 3 minutes using a pulled glass micropipette. The pipette was left in place for 2 additional minutes to minimize reflux.

Mice were treated twice daily with either FK-506 (10 mg/kg, i.p.) or cyclosporine A (10 mg/kg, i.p.) for two weeks post-transplantation as well as one day before cell grafting. At endpoint, animals were perfused with PBS followed by 4% PFA. Spinal cords were cryoprotected in 30% sucrose, embedded in OCT, and sectioned at 30 µm. Transverse sections at C2 (four segments rostral to graft site) were immunostained for GFP, and axons were quantified by blinded observers^19^.

## RESULTS

### Cyclosporine A Robustly Inhibits Neurite Outgrowth in hiPSC-Derived and Fetal Spinal Cord–Derived Neurons

To determine how clinically relevant immunosuppressants affect axon extension, we treated human iPSC-derived neurons with a panel of agents at concentrations ranging from 0.01 to 10 µg/ml. Cyclosporine A (CsA) produced a marked, dose-dependent reduction in neurite length per cell, with an 85% decrease at 10 µg/ml (**Fig. 1**). By contrast, FK-506 (tacrolimus) had no measurable inhibitory effect across the tested concentrations.

**Figure 1.**
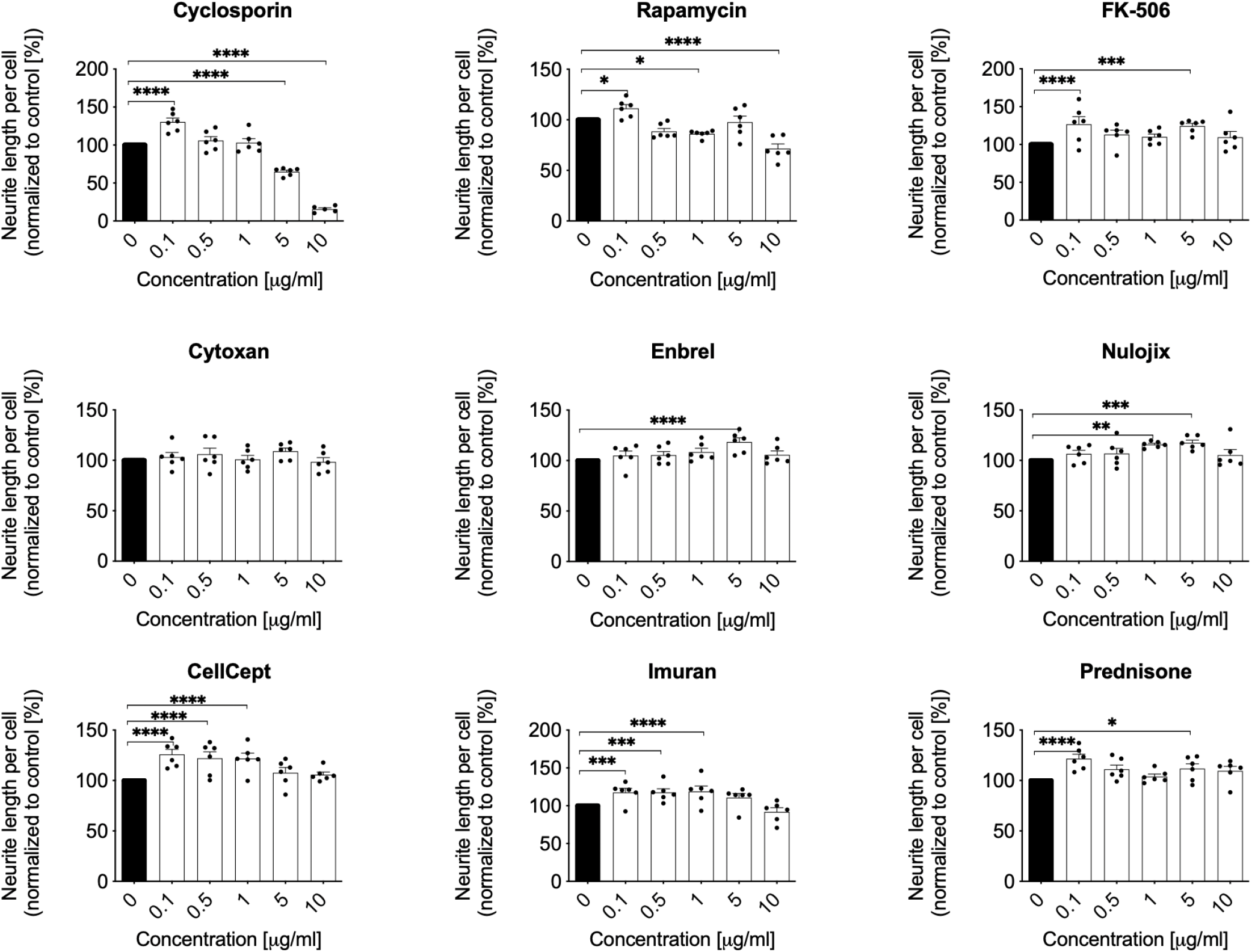
Cyclosporine Reduces Axon Growth in hiPSC-Derived Neurons In Vitro. hiPSC derived neurons were plated in 384-well plates and treated 2 hours later with immunosuppressants as indicated (final concentrations range from 0.1 to 10 µg/ml). Shown are total neurite length per cell (Mean+SEM). Cyclosporine showed a significant dose dependent reduction of neurite growth with a maximum reduction of 85% at 10 µg/ml. (****p<0.0001, one-way ANOVA, with *p<0.05, **p<0.01, ****p<0.0001 post-hoc Tukey’s test; n=2 individual experiments, n=3 wells per condition).

Other agents demonstrated variable effects. Notably, azathioprine (Imuran) significantly reduced neurite outgrowth at all tested doses, and belatacept (Nulojix) and mycophenolate mofetil (CellCept) each produced significant reductions at 10 µg/ml. These effects were statistically significant but less pronounced than the inhibition observed with CsA.

To validate these findings, we repeated the screen in an independent hiPSC-derived neuronal line. Once again, CsA produced a consistent, dose-dependent inhibitory effect, reducing outgrowth by approximately 50% at 10 µg/ml. Additionally, rapamycin significantly reduced neurite extension by ∼40% at its highest concentration (**Fig. S1**). These data indicate that while CsA exerts the most potent and consistent inhibitory effect, Imuran, Nulojix, CellCept, and rapamycin also negatively affect neurite outgrowth under some conditions.

### To evaluate whether these results extend across neural lineages, we assessed neurite outgrowth in human fetal spinal cord–derived NPCs exposed to the same drug panel

Again, CsA significantly inhibited neurite extension, with a 40–50% reduction at 5 and 10 µg/ml (**Fig. 2**). Imuran caused a similarly broad and significant suppression across doses. Nulojix and CellCept also showed significant inhibitory effects at higher doses. In contrast, FK-506, prednisone, and etanercept did not significantly alter neurite growth in this system.

**Figure 2.**
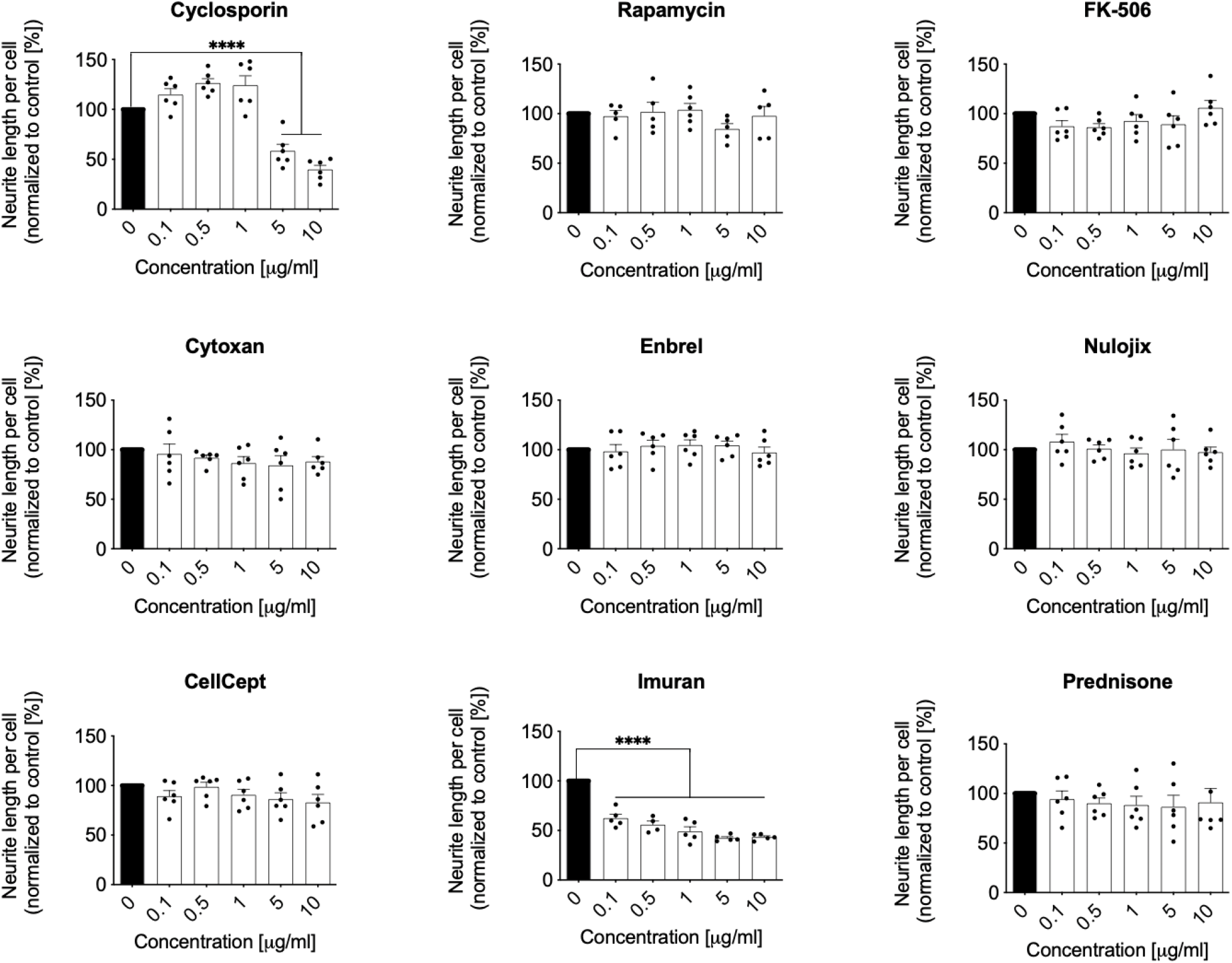
Axon Growth of Human Fetal Spinal Cord Derived NPCs is Reduced by Cyclosporine in Vitro. Human Fetal Spinal Cord Derived NPCs were plated in 96-well plates and treated 2 hours later with immunosuppressants as indicated (final concentrations range from 0.1 to 10 µg/ml). Shown are total neurite length per cell (Mean+SEM). Imuran showed an overall reduction of neurite growth at all concentrations; n=2 individual experiments, n=2-3 wells per condition). Cyclosporine showed a significant reduction of neurite growth of 40-50% at 5 µg/ml and 10 µg/ml (****p<0.0001, one-way ANOVA, with ****p<0.0001 post-hoc Tukey’s test; n=2 individual experiments, n=2-3 wells per condition).

These findings confirm that CsA consistently impairs axon extension across both stem cell– derived and primary human neural cell types, while several other immunosuppressants— particularly Imuran—can exert significant inhibitory effects under select conditions.

### DMSO Vehicle Control Does Not Affect Axonal Outgrowth

Since several tested immunosuppressants were dissolved in DMSO, we conducted a vehicle control experiment to assess whether DMSO alone alters neurite outgrowth. hiPSC-derived neurons were treated with increasing DMSO concentrations from 0.00005 to 0.50 µl/ml. No significant effect on neurite length per cell was observed at any dose tested (**Fig. S3**), confirming that the inhibitory effects observed are attributable to the immunosuppressants themselves and not the vehicle.

### Cyclosporine A Suppresses Graft-Derived Axonal Extension In Vivo, Whereas FK-506 Preserves Outgrowth

We next tested whether the in vitro effects of CsA and FK-506 translated to the in vivo setting. Human fetal spinal cord–derived NPCs expressing GFP were grafted into the intact C5 spinal cord of immunosuppressed nude mice and treated with either FK-506 or CsA. Two weeks later, axon extension into the rostral host spinal cord was quantified by immunostaining for GFP. Mice treated with FK-506 exhibited robust graft-derived axon extension into the white matter three segments rostral to the graft site. In contrast, CsA-treated mice showed a dramatic reduction in axonal growth (**Fig. 3A**).

**Figure 3.**
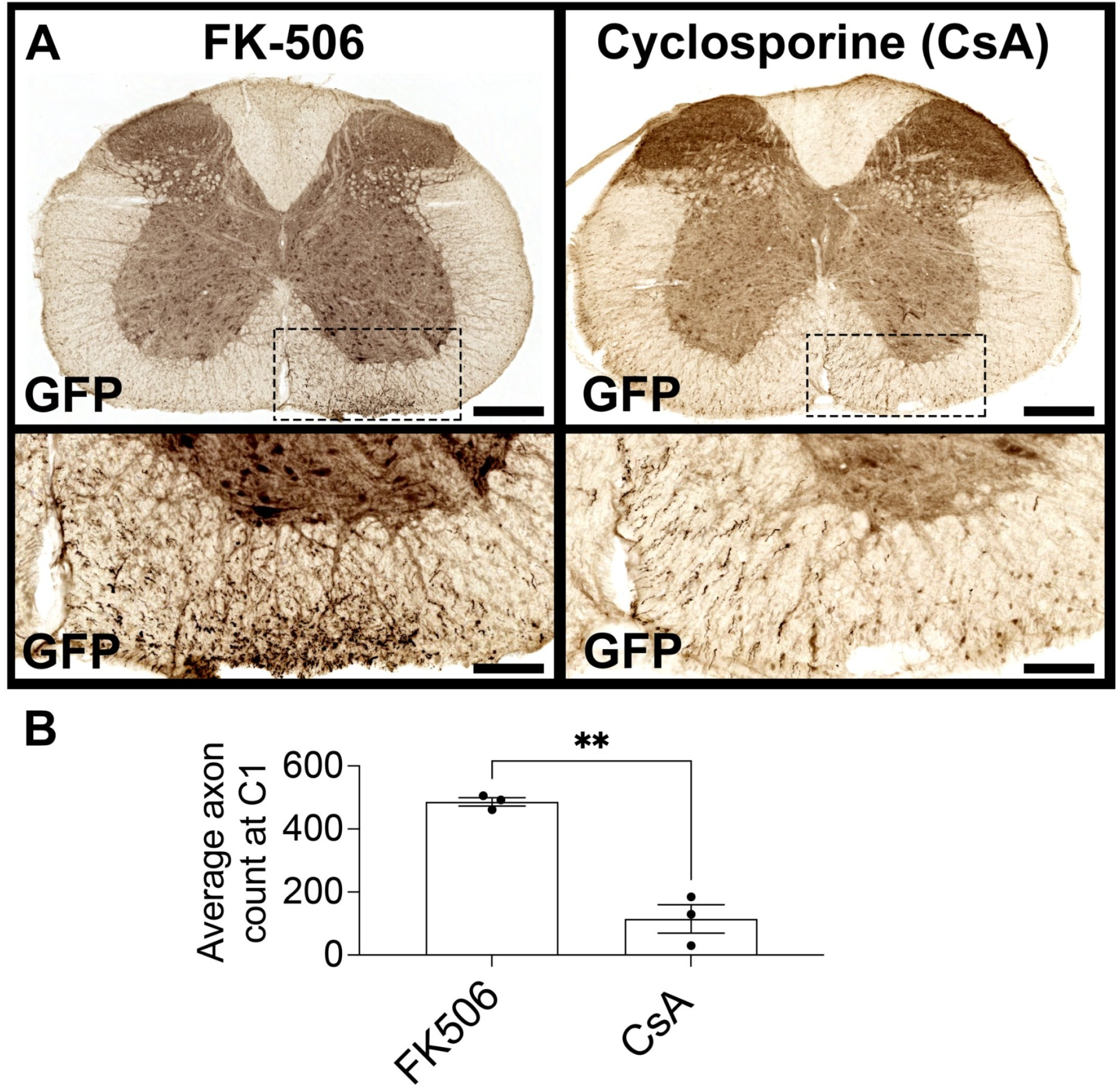
Human Fetal Spinal Cord Derived NPCs Extend Reduced Numbers of Axons into Spinal Cord White Matter 3 Segments Above Graft Site in Response to Cyclosporine Treatment. **(A)** GFP-expressing fetal human spinal cord-derived NPCs were grafted into sites of C5 intact spinal cord. Mice were either treated daily with FK-506 (left panel) or Cyclosporine (right panel). Two weeks after grafting, transverse sections of spinal cord were immunolabeled for GFP (black). NPC graft-derived axons in transverse sections of mouse spinal cord treated with Cyclosporine demonstrates reduced numbers of axons extending into rostral host spinal cord white matter, 3 spinal segments above the graft site. **(B)** Quantification of data in panels A. Mean+SEM (**p<0.01, un-paired, two-tailed t-test; n=3 mice per group). Scale bar: A, 500μm; magnification, 25μm.

Quantification revealed significantly fewer GFP+ axons at the C2 level in the CsA group (**Fig. 3B; *p* < 0.01**). This effect was consistent across all animals. Images of transverse sections from all biological replicates confirmed this pattern: axon density was substantially lower in CsA-treated mice, while FK-506–treated mice consistently exhibited axon-rich regions in the dorsal and lateral columns (Figure S2).

## DISCUSSION

This study demonstrates that among multiple clinically relevant immunosuppressants, only cyclosporine A (CsA) significantly impairs axon elongation in human neurons, both in vitro and in vivo. Using two distinct human cell models—hiPSC-derived neurons and fetal spinal cord–derived NPCs—we show that CsA suppresses neurite outgrowth in a dose-dependent manner, with no comparable inhibitory effects observed for FK-506, rapamycin, or other tested agents. These findings were reinforced in an in vivo model of spinal cord grafting, where mice treated with CsA exhibited markedly reduced extension of graft-derived axons into the host spinal cord compared to animals treated with FK-506. To our knowledge, this is the first direct comparison of axon-regenerative outcomes under different immunosuppressive regimens in the context of human neural grafts.

Our results are consistent with earlier observations that CsA can affect neuronal differentiation and growth. CsA inhibits calcineurin, a calcium/calmodulin-dependent phosphatase involved in diverse neuronal functions, including neurite extension, synaptic plasticity, and axonal guidance. Inhibition of calcineurin has been shown to modulate the activity of NFAT and other downstream targets critical for cytoskeletal remodeling and axonal pathfinding. Although calcineurin inhibition is common to both CsA and FK-506, prior studies suggest they engage different protein targets—cyclophilin A and FKBP12, respectively—which may lead to divergent downstream effects in neurons^20^ (Matsuda & Koyasu, 2000). The stark contrast in axon outgrowth between CsA- and FK-506–treated neurons observed here supports the view that CsA’s inhibitory effects may be mediated through off-target interactions or differential modulation of calcineurin isoforms in neural cells.

The lack of a significant effect from rapamycin, mycophenolate, prednisone, or azathioprine suggests that not all immunosuppressants interfere with neurite extension at clinically relevant doses. Rapamycin is known to inhibit mTOR signaling, a pathway with well-established roles in axon regeneration, particularly in the peripheral nervous system and in models of CNS injury^15^. However, our results indicate that short-term rapamycin exposure at standard immunosuppressive concentrations does not impair neurite growth in human NPCs or hiPSC-derived neurons. This is in agreement with recent findings showing that mTOR inhibition may have a more pronounced effect on regeneration in mature CNS neurons than on developing or graft-derived axons.

Importantly, these findings have direct translational relevance for neural stem cell transplantation strategies in spinal cord injury (SCI). Immunosuppressive therapy is routinely administered in clinical and preclinical stem cell transplantation to prevent immune-mediated rejection, even when autologous or iPSC-derived grafts are used^10^. Our data underscore that the choice of immunosuppressive agent is not merely a matter of immunological compatibility but also of neuroregenerative permissiveness. In this context, FK-506 emerges as a preferred agent due to its lack of negative impact on axon outgrowth and its consistent use in recent SCI clinical trials^7^.

Beyond selecting agents with minimal adverse effects, there is also growing interest in identifying immunosuppressants that may actively enhance neural regeneration. FK-506 has previously been shown to exert neurotrophic effects in some contexts, including enhanced axonal regeneration following peripheral nerve injury^21^. While we did not observe enhanced growth beyond baseline in FK-506–treated neurons, it is possible that in longer-term or injury models, its regenerative potential may be more evident. Future studies could explore combination therapies that optimize both immune tolerance and axon regeneration.

### CONCLUSION

In summary, our findings highlight a previously underappreciated consequence of immunosuppressant selection in neural graft protocols: CsA significantly impairs axonal outgrowth from human neurons, whereas FK-506 preserves regenerative potential. These results argue for rigorous preclinical assessment of immunosuppressive regimens not only in terms of graft survival but also in their compatibility with neuronal growth and integration.

Optimizing immunosuppression will be a critical determinant of success in future cell-based therapies for SCI and other neurodegenerative disorders.

## Abbreviations

SCI: Spinal cord injury
NSC: Neural stem cell
NPC: Neural progenitor cell
hiPSC: Human Induced pluripotent stem cell
CsA: Cyclosporine A
FK-506: Tacrolimus
GFP: Green fluorescent protein
PDL: Poly-D-lysine
IL-2: Interleukin-2
mTOR: Mechanistic target of rapamycin

## FIGURE CAPTIONS

**Figure S1. Axon Growth of hiPSC Derived Neurons is Reduced by Cyclosporine in Vitro (Second patient derived IPSC line)**

hiPSC derived neurons were plated in 384-well plates and treated 2 hours later with immunosuppressants as indicated (final concentrations range from 0.1 to 10 µg/ml). Shown are total neurite length per cell (Mean+SEM). Cyclosporine showed a significant 50% reduction of neurite growth at 10 µg/ml. Rapamycin showed a significant 40% reduction of neurite growth only at 10 µg/ml (****p<0.0001, one-way ANOVA, with *p<0.05, post-hoc Tukey’s test; n=2 individual experiments, n=3 wells per condition),

**Figure S2. Human Fetal Spinal Cord Derived NPCs Extend Reduced Numbers of Axons into Spinal Cord White Matter 3 Segments Above Graft Site in Response to Cyclosporine Treatment**

GFP-expressing fetal human spinal cord-derived NPCs were grafted into sites of C5 intact spinal cord. Mice were either treated daily with FK-506 (left panel) or Cyclosporine (right panel). Two weeks after grafting, transverse sections of spinal cord were immunolabeled for GFP (black). NPC graft-derived axons in transverse sections of mouse spinal cord treated with cyclosporine demonstrates reduced numbers of axons extending into rostral host spinal cord white matter, 3 spinal segments above the graft site. Shown are example images of all biological replicates corresponding to Figure 3A. Scale bar: A, 500μm; magnification, 25μm.

**Figure S3. DMSO Vehicle Does Not Affect Neurite Outgrowth in hiPSC-Derived Neurons.** Human iPSC-derived neurons were plated in 384-well plates and treated 2 hours later with DMSO at increasing concentrations (0.00005 to 0.50 µl/ml). Total neurite length per cell was quantified via automated high-content imaging 24 hours later using the MetaXpress neurite outgrowth module. No significant changes in neurite outgrowth were observed across the tested concentrations, indicating that DMSO alone does not impair axonal growth. Data are presented as mean ± SEM (n = 2 independent experiments, n = 3–6 wells per condition).

## ACKNOWLEDGMENTS

This work was supported by NIH R01 NS104442, the Veterans Administration (B7332R), and The Dr. Miriam and Sheldon G. Adelson Medical Research Foundation (to M.H.T.).

