## Supplementary figures and images for "Clinical Relevant Immunosuppressive Drugs Differentially Modulate Axonal Outgrowth from Human Stem Cell–Derived Neurons"

### Supplemental Figure 1

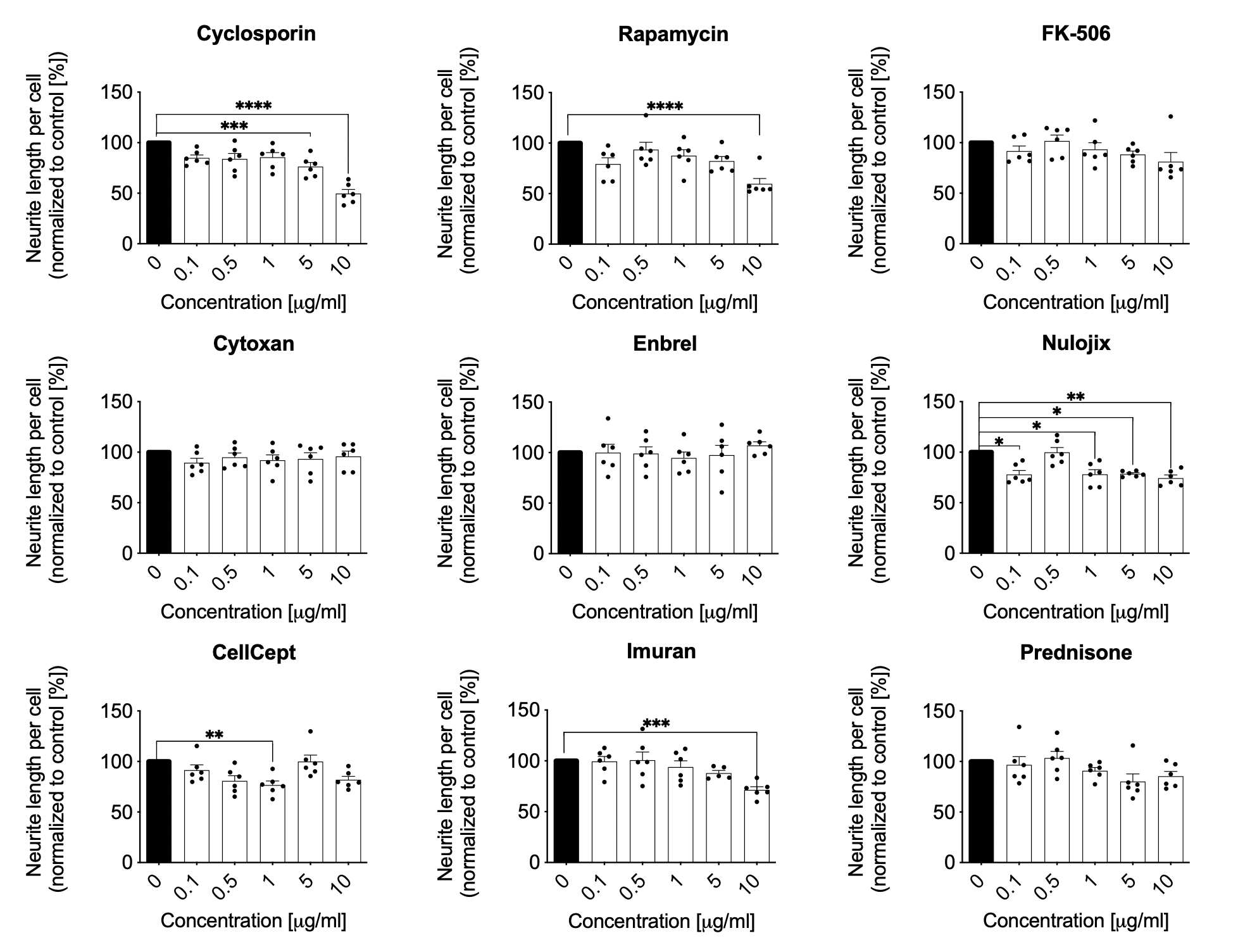

### Supplemental Figure 2

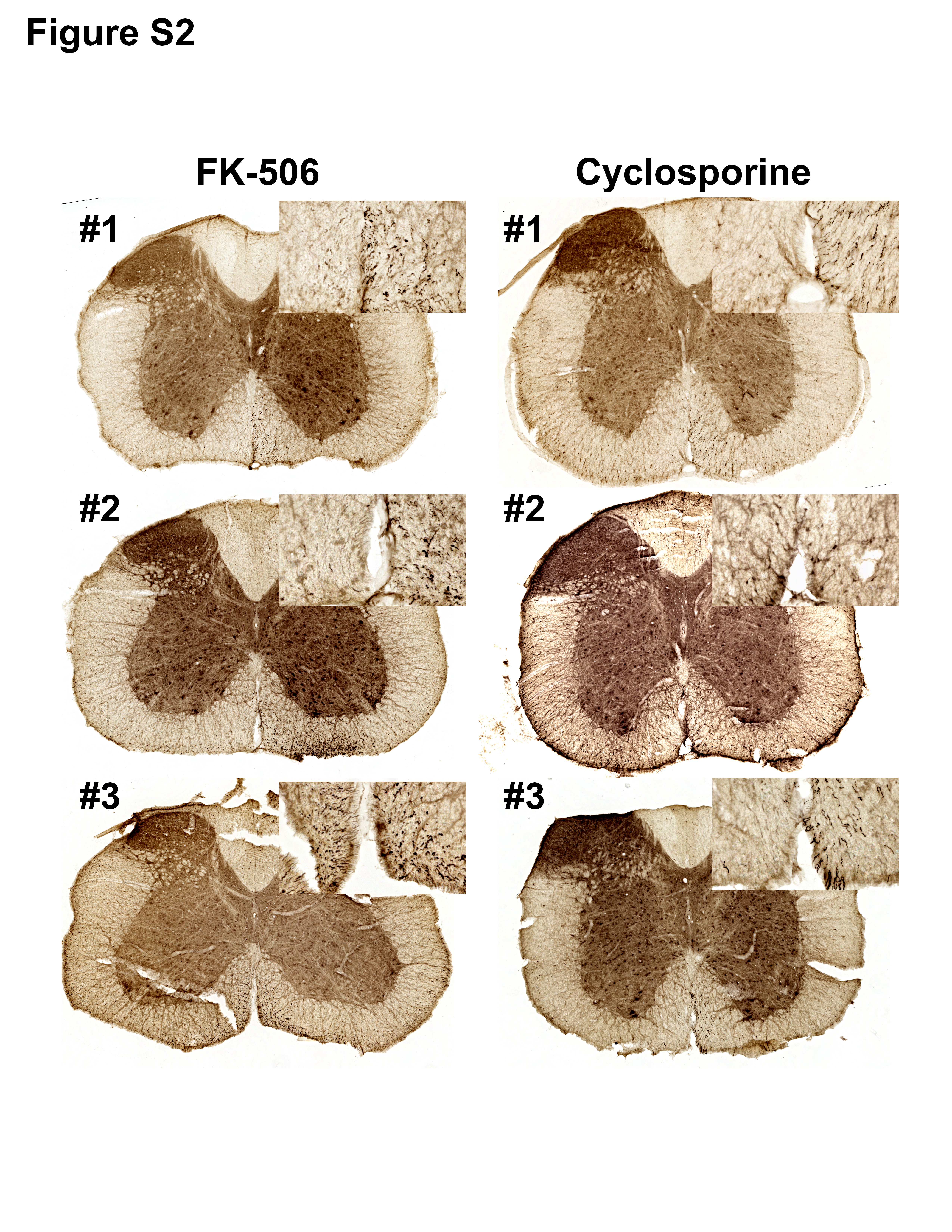

### Supplemental Figure 3

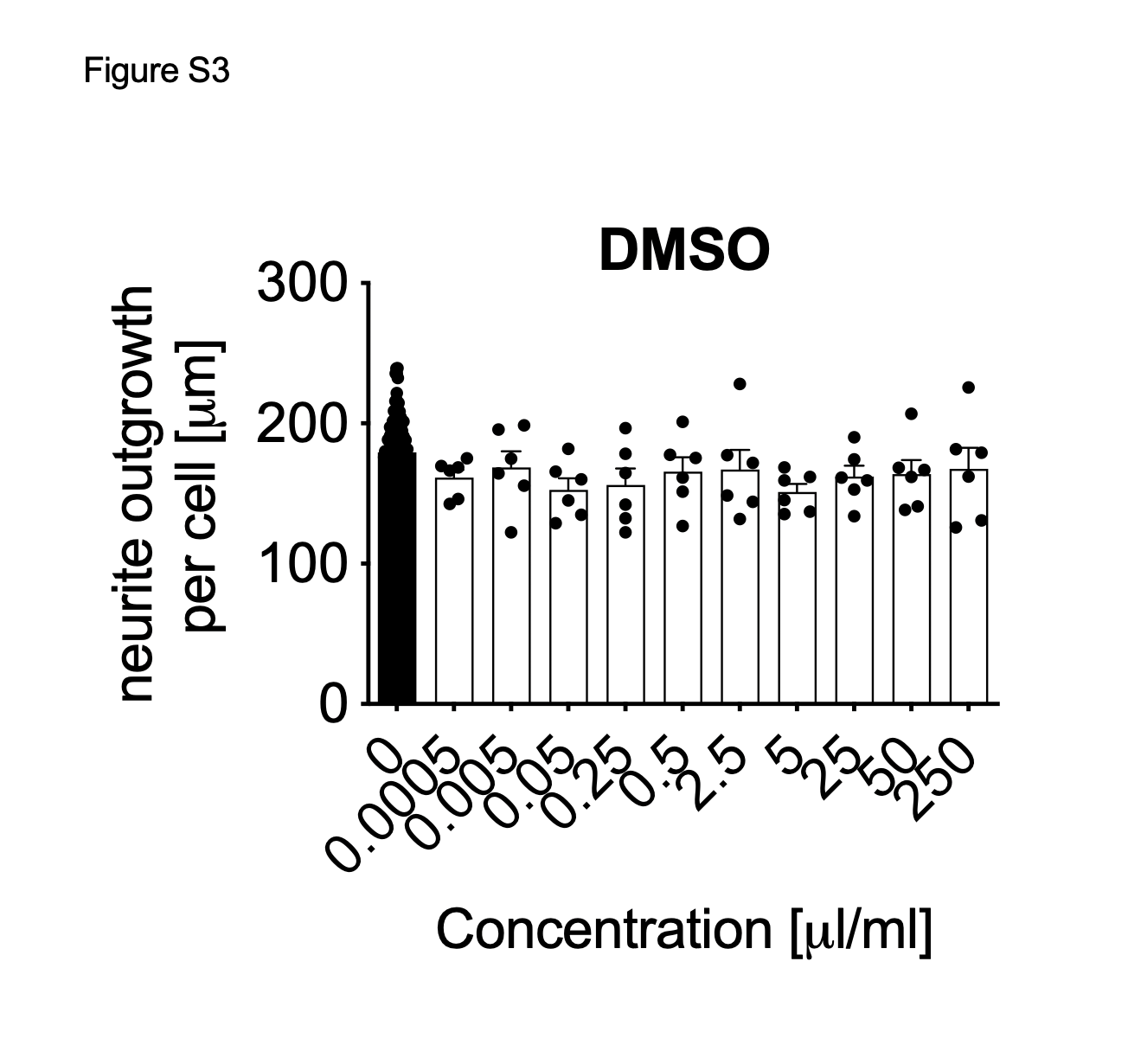
